# NG2-Glia Heterogeneity Across Cortical Layers

**DOI:** 10.1101/2025.07.23.666263

**Authors:** Sonsoles Barriola, Lina María Delgado-García, Ana Cristina Ojalvo-Sanz, Carolina Pernia Solanilla, María Figueres-Oñate, Laura López-Mascaraque

## Abstract

NG2-glia are a unique and heterogeneous glial cell population with diverse roles in the central nervous system. However, their morphological diversity across brain regions and cortical layers remains poorly understood. Here, we use StarTrack labeling and in utero electroporation at embryonic day 14 (E14) to reconstruct individual NG2-glial cells in the adult mouse cortex and corpus callosum. Through detailed two- and three-dimensional morphometric analyses, including Sholl analysis, principal component analysis, and hierarchical clustering, we uncover striking layer-specific patterns. NG2-glia in deep cortical layers (L5–6) exhibit significantly larger somatic areas, more elaborate arborizations, and higher process complexity compared to those in superficial layers (L1–4) and the corpus callosum. In contrast, NG2-glia in layer 1 and the corpus callosum share a compact morphology characterized by smaller somata and simplified processes, suggesting common microenvironmental constraints. Moreover, Sholl analysis, principal component analysis, and hierarchical clustering reveal distinct morphological subpopulations within the NG2-glial population and highlight heterogeneity in upper cortical layers. Comparative analyses with astrocytes reveal fundamental structural differences: NG2-glia have thinner, longer processes and larger enclosing radii but occupy smaller volumes, whereas astrocytes form denser, more compact arbors with higher branch numbers.

Together, our finding establish the first comprehensive morphological atlas of cortical adult NG2-glia, highlighting region- and layer-specific adaptations that likely underlie their diverse roles in CNS physiology and repair.

**Graphical abstract:** 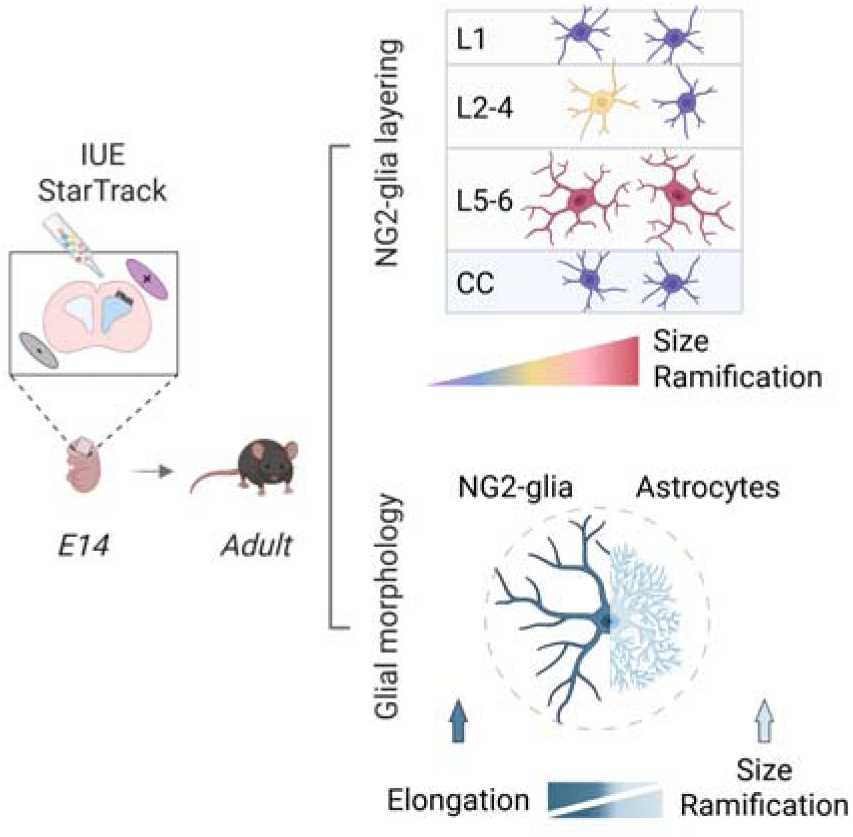

**Main points:**

- Morphological variation of NG2-glial cells by cortical layers
- Cells in deep cortical layers are larger and more complex
- NG2-glia in upper layers and corpus callosum share similar morphology
- NG2-glia vs. astrocytes: distinct structural features

## 1 Introduction

The glial landscape of the mammalian central nervous system (CNS) encompasses a diverse array of cell types that support neuronal function and maintain homeostasis. Among these, NG2-glia have emerged as a distinct population within the central nervous system, differing significantly from other neural cell types such as neurons, oligodendrocytes, astrocytes, and microglia (Butt et al., 2005; Nishiyama et al., 2009; Richardson et al., 2011; García-Marqués et al., 2014; Dimou and Gallo, 2015; Figueres-Onãte et al., 2016; Sánchez-González et al., 2020). Though often referred to as oligodendrocyte progenitor cells (OPCs) due to their capacity to differentiate into mature oligodendrocytes (Nishiyama et al., 1999), recent studies reveal their significant heterogeneity during both development and adulthood (Viganò et al., 2016; Spitzer et al., 2019; Marisca et al., 2020; Ojalvo-Sanz et al., 2025). This growing body of research highlights NG2-glia as a specialized glial population with unique characteristics and functions, setting them apart from other macroglial cells in the brain.

A defining feature of NG2-glia is their remarkable capacity for proliferation and differentiation potential throughout life, particularly in response to injury (Simon et al., 2011). These progenitor-like properties are coupled with unique functional traits that are not observed in other glial populations. For instance, NG2-glia establish functional synaptic-like connections with neurons in the adult brain (Bergles et al., 2000; Lin and Bergles, 2003; Ge et al., 2006), characteristic not typically seen other glial cells. This synaptic interaction indicates that NG2-glia play an active role in neuronal communication and network plasticity. Furthermore, NG2-glia participate in the clearance of neuronal elements (Auguste et al., 2022; Buchanan et al., 2022), a process that resembles microglial phagocytosis but with distinct functional implications. This dual role, both synaptic and phagocytic, underscores the functional versatility of NG2-glial cells.

Similar to astrocytes, NG2-glia exhibit regional differences in their interactions with neurons, as well as in their electrophysiological properties, particularly between white and gray matter (Chittajallu et al., 2004; Serwanski et al., 2017). Astrocyte morphological diversity is associated with neuronal connectivity across cortical layers and brain regions, and is linked to functional differences (Xin Tan et al., 2023). NG2-glia appear to follow a similar pattern, as they undergo hypertrophy following brain injury, indicating a dynamic interplay between their structural features and functional state (Simon et al., 2011; Honsa et al., 2012; Bonfanti et al., 2017; Coppolino et al., 2018; Barriola et al., 2020). However, despite these insights, a comprehensive morphological characterization of NG2-glia is lacking, limiting our understanding of their potential heterogeneity, a key factor in their diverse roles within the adult brain.

In this study, we used the StarTrack labeling via in utero electroporation at embryonic day 14 to achieve sparse, multicolor cytoplasmic marking of NG2-glia in adult mice. We reconstruct single cells from cortical layers 1–6 and the corpus callosum, applying Sholl analysis, principal component analysis, and hierarchical clustering to uncover layer- and region-specific morphologies. By contrasting NG2-glia with adjacent astrocytes, we delineate unique structural strategies, laying a foundation for understanding how morphological heterogeneity relates to the varied functions of NG2-glia in the adult CNS.

## 2 Material and Methods

### 2.1 Animals

C57BL/6 mice were hosed and maintained at the animal facility of the Cajal Institute under standard conditions. All animal procedures were conducted in accordance with the European Directive 2010/63/EU and the regulations established by the Spanish Animal Care and Use Committee. Experimental protocols received approval from the Bioethics Committee of the Community of Madrid (Reference: PROEX 314/19).

### 2.2 StarTrack DNA Plasmids

Adult NG2-glial cells were labelled using modifications of the StarTrack technique: the UbC-(GFAP-PB)-StarTrack and UbC-StarTrack to follow the entire progeny of targeted neural progenitor cells (NPCs) (Figueres-Onãte et al., 2016; Ojalvo-Sanz et al., 2024). These strategies allowed the identification of NG2-glia in the adult mouse brain.

Plasmids were prepared and purified using the NucleoBond Xtra Maxi kit (Macherey-Nagel, France, #740414.50). Competent DH5α E. coli cells were transformed with the desired target plasmid via heat shock, plated on agar supplemented with ampicillin and cultured overnight. A single resistant colony was expanded in LB broth containing ampicillin (10 µg/mL). After bacterial growth, the DNA plasmid was extracted via column-based purification.

### 2.3 In Utero Electroporation

In utero electroporation (IUE) was performed at embryonic day 14 (E14) as previously described. The UbC-StarTrack or UbC-(GFAP-PB)-StarTrack mix was injected into the lateral ventricles of each embryo, followed by the delivery of five electric pulses (35 mV, 50 ms duration) of 950 ms intervals using an electroporator. For selective targeting of dorsal pallial progenitors, the anode was positioned over the dorsal part of the embryo head. After electroporation, the uterine horns were carefully repositioned into the abdominal cavity, and the embryos were allowed to develop until the designated experimental stage.

### 2.4 Tamoxifen Administration

To activate Cre-LoxP recombination and eliminate non-integrated plasmids, tamoxifen (5 mg/40 g, Sigma-Aldrich) was administered at postnatal stage P7. Tamoxifen was dissolved in corn oil at a concentration of 20 mg/ml in corn oil, and delivered via intraperitoneal injection.

### 2.5 Immunohistochemistry

Mice were anesthetized and perfused transcardially with 4% paraformaldehyde (PFA). Brains were post-fixed for an additional hour, and coronal sections (50 µm) were obtained using a vibratome. Sections were washed and permeabilized three times in PBS containing 0.5% Triton X-100 (PBS-T), followed by additional washes in f 0.1% PBS-T. For PDGFR_α_ immunolabeling (denoted by an asterisk in Table 1), an additional step 20-minute treatment in 10% methanol was applied.

**Table 1.** List of primary and secondary antibodies.

| Antibody | Host | Use | Source | Cell Type |
| --- | --- | --- | --- | --- |
| CTIP2 | Rat | 1:100 | Abcam | Neurons in deeper layers |
| Cux1 | Rabbit | 1:300 | Proteintech | Neurons in upper layers |
| PDGFRα* | Rabbit | 1:300 | Cell Signaling | NG2-glial cells |
| Alexa Fluor 633 anti-Rabbit | Goat | 1:1000 | ThermoFisher | Secondary antibody |
| Alexa Fluor 647 anti-Rat | Goat | 1:1000 | Invitrogen | Secondary antibody |

Sections were subsequently incubated at room temperature for at least 2 hours in a blocking solution (5% normal goat serum (NGS) in 0.1% PBS-T), followed by overnight incubation at 4 °C with primary antibodies (listed in Table 1) diluted in the same blocking solution. The next day, sections were washed in 0.1% PBS-T and incubated for 2 hours at room temperature with the corresponding secondary antibodies. Final washes were performed with 0.1% PBS-T and PBS1X, and sections were mounted on glass slides and coverslipped using Mowiol (Polysciences, #17951).

### 2.6 Image Acquisition

Confocal imaging was performed using Leica Stellaris-8 (Germany). For clonal analysis, the six StarTrack fluorescent reporters and the immunohistochemical signals were acquired with distinct spectral channels, ensuring no overlap (see Table 2).

**Table 2.**

| Stellaris-8 |  | mT-Sapphire | mCerulean | EGFP | YFP | mKO | mCherry |
| --- | --- | --- | --- | --- | --- | --- | --- |
|  | Excitation | 405 | 440 | 489 | 522 | 548 | 587 |
|  | Emission | 525-540 | 455-490 | 497-510 | 529-544 | 556-580 | 597-625 |
|  | Intensity | 6.5% | 8.01% | 2,1% | 4% | 2% | 8.01% |

### 2.7 Morphometric Analysis

A total of 36 NG2-glial cells were analyzed from the cortex and corpus callosum of three mice electroporated at embryonic day 14 (E14) with StarTrack plasmids. Reconstructions were performed manually using the Simple Neurite Tracer (SNT) plugin in Fiji, tracing the entire arborization of each cell from the soma through the confocal Z-stacks (1024 × 1024 resolution; 40X objective; 1 µm Z-step). After reconstruction, a 0.01 threshold was applied to fill traced structures, allowing the extraction of morphometric parameters such as area, perimeter, circularity, solidity, convex hull area, and volume. Traced structures were subsequently skeletonized to generated a 3D mask for analysis of process complexity, including branch number and maximum branch length. Sholl analysis was performed based on the SNT-derived data.

### 2.8 Data and Statistical Analyses

Data analysis and visualization were conducted using R version 4.1.2 (R Core Team, 2021) and GraphPad Prism 8. The normality was assessed using the Kolmogorov-Smirnov test. For normally distributed datasets, unpaired two-tailed Student’s *t*-tests were used for two-group comparisons, and one-way ANOVA for comparisons involving more than two groups. For non-normally distributed data, Kruskal-Wallis tests followed by Dunnett’s post hoc test for multiple group comparisons or Mann-Whitney tests (a non-parametric two-tailed unpaired *t*-test), for two-group comparisons were applied. A p-value below 0.05 was considered statistically significant (95% confidence interval). Statistically significant differences are indicated in figures with asterisks as follows: * p < 0.05, ** p < 0.01, *** p < 0.001.

## 3 Results

### 3.1 Layering of cortical NG2 Glia

To map NG2-glia morphology across cortical layers and the corpus callosum, we performed IUE at E14 (Figure 1A), using two complementary StarTrack strategies: the UbC-StarTrack and the UbC-(GFAP-PB)-StarTrack (Figure 1B). In both approaches, fluorescent proteins were expressed in the nucleus and/or cytoplasm under the control of the ubiquitous Ubc promoter. To selectively remove episomal constructs, tamoxifen was administered at P7, triggering Cre recombinase activity via the Cre-LoxP system (Figure 1A,nB). The main distinction between the two strategies lay in the regulation of the PiggyBac transposase (hyPBase): the CMV promoter drove broad expression across NPCs, whereas the GFAP promoter provided a more selective labeling of GFAP+ progenitors at the time of electroporation. NG2-glia was analyzed at postnatal days 60 and 90, stage when myelination is largely complete (Figure 1A). Both strategies successfully labeled NG2-glial cells in the cortex and corpus callosum (Figure 1B).

**Figure 1.**
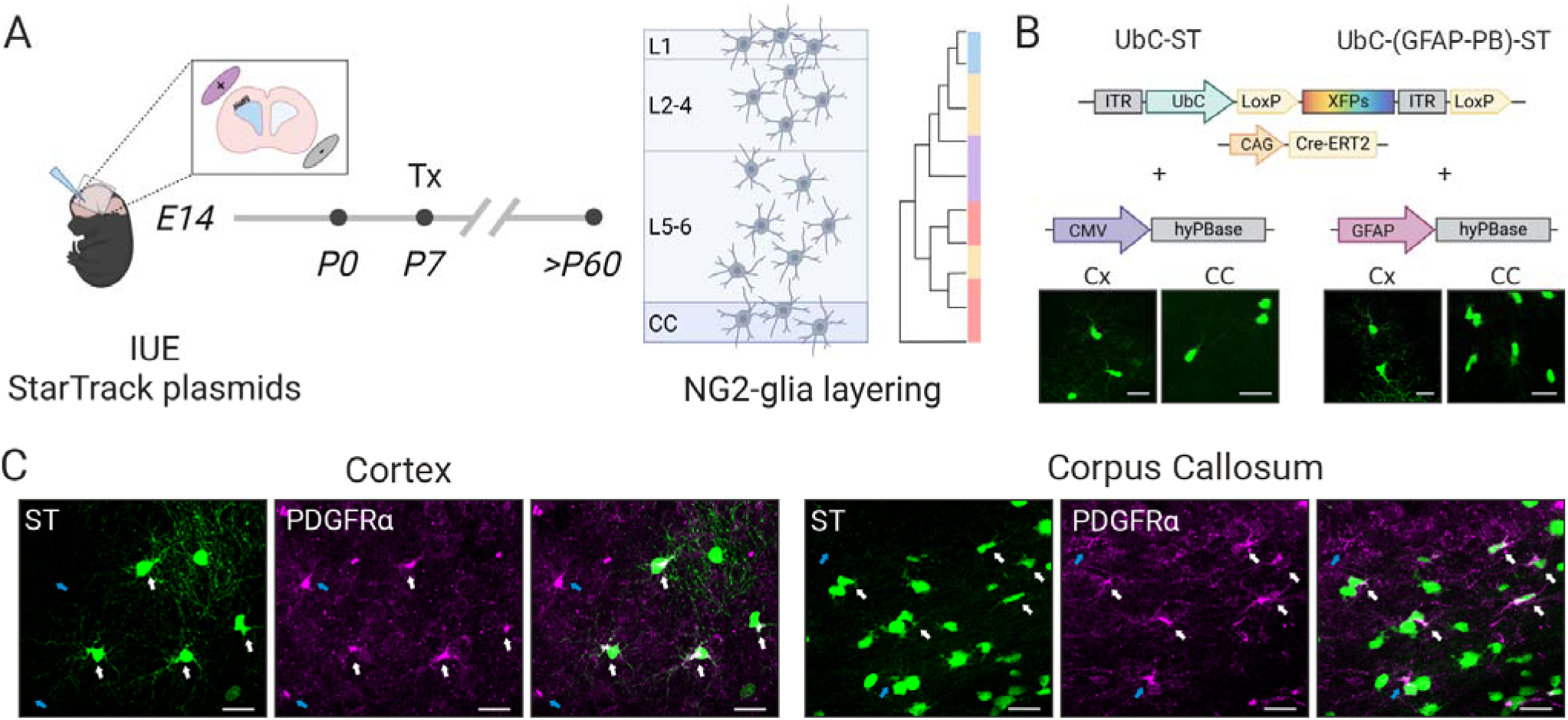
**Morphological Analysis of NG2-Glia in the Cortex and Corpus Callosum Using StarTrack.** (A) Experimental design: NG2-glia were traced using StarTrack via in utero electroporation (IUE) at embryonic day 14 (E14). Tamoxifen was administered at postnatal day 7 (P7), and NG2-glia morphology was analyzed in adult brains, specifically in the cortex (Cx) and corpus callosum (CC). (B) Two distinct StarTrack strategies were employed. In both the ubiquitous UbC promoter drove the expression of the fluorescent reporters, while the PiggyBac transposase (hyPBase) was regulated either by the CMV promoter (for broad NPC targeting) or the GFAP promoter (for selective targeting of GFAP+ NPCs). Both strategies successfully labeled NG2-glial cells in the cortex (Cx) and corpus callosum (CC), enabling detailed visualization of their morphology. (C) StarTrack[NG2-glia. White arrows indicate NG2-glial cells colocalizing with PDGFRα], while blue arrows indicate PDGFRα[cells that are not StarTrack]. No major morphological differences were detected across labeling strategies. Scale bars represent 20 μm.

A key advantage of StarTrack system is its ability to label the cytoplasm, allowing high-resolution visualization of cell morphology and sparse population labeling (Figure 1B-C). Using this approach, we analyzed 36 NG2-glial cells across cortical layers and the corpus callosum, identified by PDGFR_α_ immunostaining (Figure 1C).

### 3.2 Quantitative Assessment of NG2-Glia Morphology

#### Two-Dimensional Morphometric Parameters of NG2-Glial Cells

Morphological reconstruction of NG2-glia was performed using the Simple Neurite Tracer (SNT) plugin available in FIJI. This semi-automated tool enabled manual tracing of cellular branches starting from the soma and extending through various z-axis planes. SNT enables extraction of both two-dimensional (2D, Figure 2A, B) and three-dimensional (3D, Figure 2C, D) morphological features.

**Figure 2.**
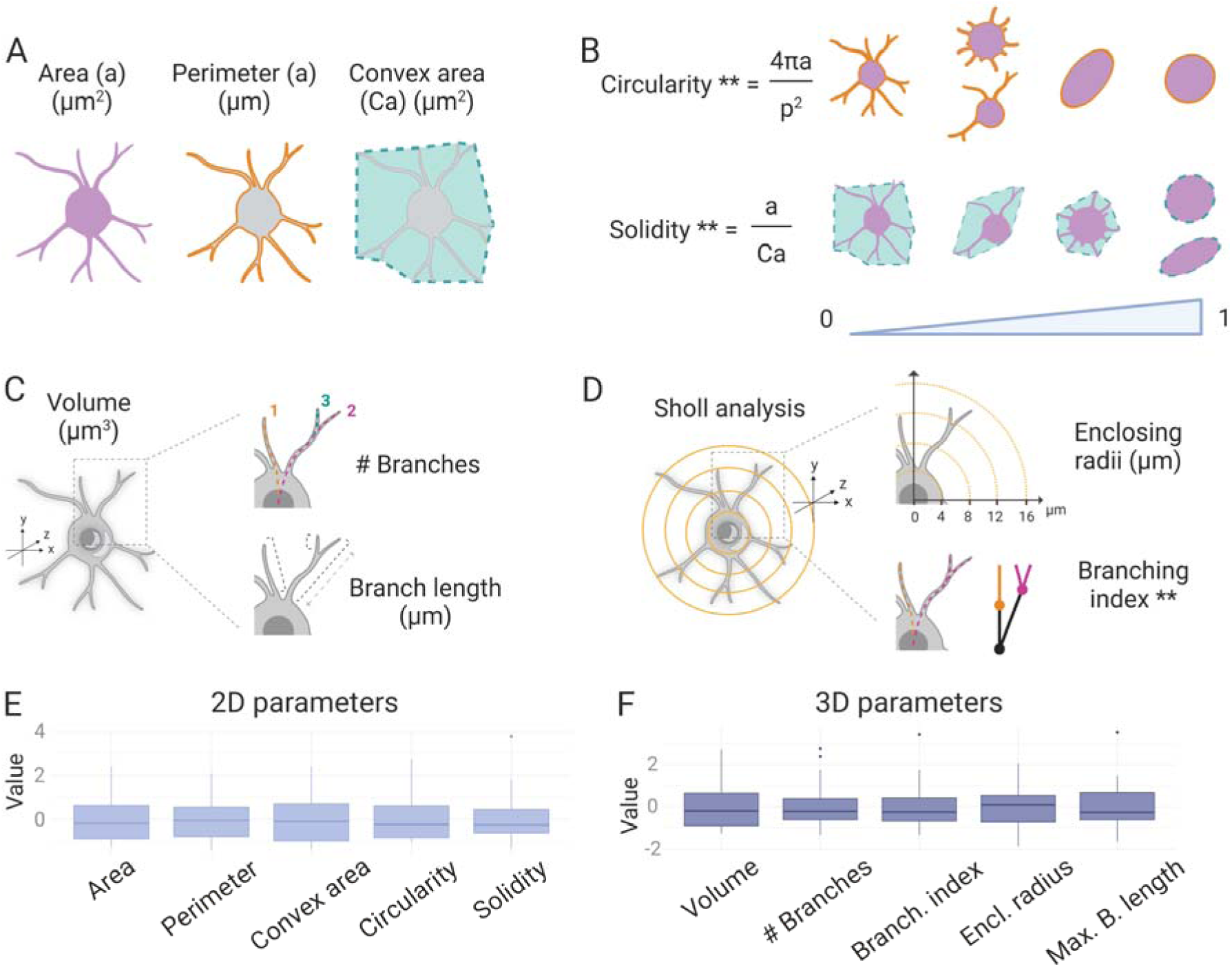
Morphometric Analysis of NG2-Glial Cells Using Two-Dimensional and Three-Dimensional Parameters. (A, B) Two-Dimensional Morphometric Parameters: (A) Size-related features including area (µm²), perimeter (µm), and convex hull area (µm²). (B) Shape descriptors circularity and solidity. (C, D) Three-Dimensional Morphometric Parameters: (C) Volume (µm³), total number of branches, and maximum branch length (µm). (D) Sholl analysis results showing process complexity and enclosing radii. (E, F) Population Variability: Box plots depicting the distribution of 2D features (E) and 3D features (F) across the NG2-glial population. Z-score normalization was applied to allow cross-cell comparisons. All morphometric data were normalized using Z-scores to allow comparison across individual cells.

For 2D analyses we quantified size descriptors such as area (µm²), perimeter (µm), and convex hull area (µm²) offer insights into cell size and structure (Figure 2A). The area reflects the total space covered by the cell, while the perimeter indicates the complete length around the cell’s outline. The convex hull area describes the area of the smallest convex shape that fully contains the cell. Furthermore, shape descriptors like circularity and solidity help define the geometry of the cell (Figure 2B). Circularity assesses how similar the cell shape is to a perfect circle and is computed as:

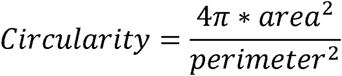

A circularity score of 1 represents a perfectly round object, whereas lower scores suggest more elongated or irregular forms.

Solidity measures how compact the cell is and is defined as the proportion of the area relative to its convex hull area:

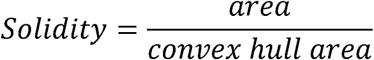

Values approaching 1 represent more compact shapes, whereas lower values suggest the presence of concave regions.

#### Three-Dimensional Characterization and Sholl Analysis

For 3D analysis of NG2-glia (Figure 2C, D), we assessed structural and size-related features, including volume (µm³), total branch count, and the length of the longest branch (Figure 2C). Volume represents the overall three-dimensional space taken up by the cell, while the number of branches indicates how many processes emerge from the soma, serving as a proxy for the cell’s morphological complexity. The maximum branch length corresponds to the longest individual extension.

Following the acquisition of arborization data through SNT analysis, we performed a Sholl analysis (Figure 2D), a technique used to evaluate the distribution and complexity of cellular branching. This approach consists of placing concentric spheres centered on the cell body, increasing the radius at fixed steps (4 µm in this case). We quantified the number of times the cell’s branches intersected with each sphere, generating a distance-dependent measure of NG2-glia arbor complexity.

From this analysis, we extracted essential 3D metrics, including the enclosed radii (µm) and the branching index. The enclosed radii indicate how far the branches extend from the soma, while the branching index represents the ratio between the number of sphere intersections and the total branch length, offering an estimate of arbor complexity.

This methodology enabled a detailed characterization of NG2-glia morphology in dorsal brain areas. To assess variation across the cell population, we applied z-score normalization to all parameters, ensuring comparability between cells. Both 2D metrics (Figure 2E) and 3D metrics (Figure 2F) revealed substantial heterogeneity in NG2-glia branching within cortical layers.

### 3.3 Regional Morphological Clusters of NG2-Glia

Due to the functional heterogeneity observed between NG2-glia in gray and white matter regions, we aimed to investigate whether these regional distinctions contribute to their morphological diversity. To begin, we assessed the principal factors driving this variability by applying the Multimodal Index (MMI) to each morphological parameter (Schweitzer and Renehan, 1997; Delgado-García et al., 2024). The MMI is a metric designed to determine how well a parameter can support data clustering based on its distribution pattern. It incorporates two statistical moments—skewness (M3) and kurtosis (M4)—offering a detailed assessment of the shape characteristics of each parameter’s distribution.

The MMI is calculated as follows:

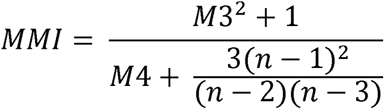

Skewness evaluates the asymmetry of a dataset’s distribution, while kurtosis reflects how much the distribution diverges from a normal distribution.

To identify the most relevant parameters for clustering, we employed MMI. In line with previous research, we considered parameters with MMI values greater than 0.55 as significant for clustering (Schweitzer and Renehan, 1997; Delgado-García et al., 2024). The most informative features defining NG2-glia clusters were circularity, solidity, number of branches, branching index, and maximum branch length (Figure 3A). These parameters were selected for Principal Component Analysis (PCA) to examine the distribution of NG2-glial cells from the cortex (n = 28) and corpus callosum (n = 8), aiming to explore potential region-specific clustering (Figure 3B). PCA enabled dimensionality reduction of the NG2-glia morphological data while preserving the variation captured by each feature.

**Figure 3.**
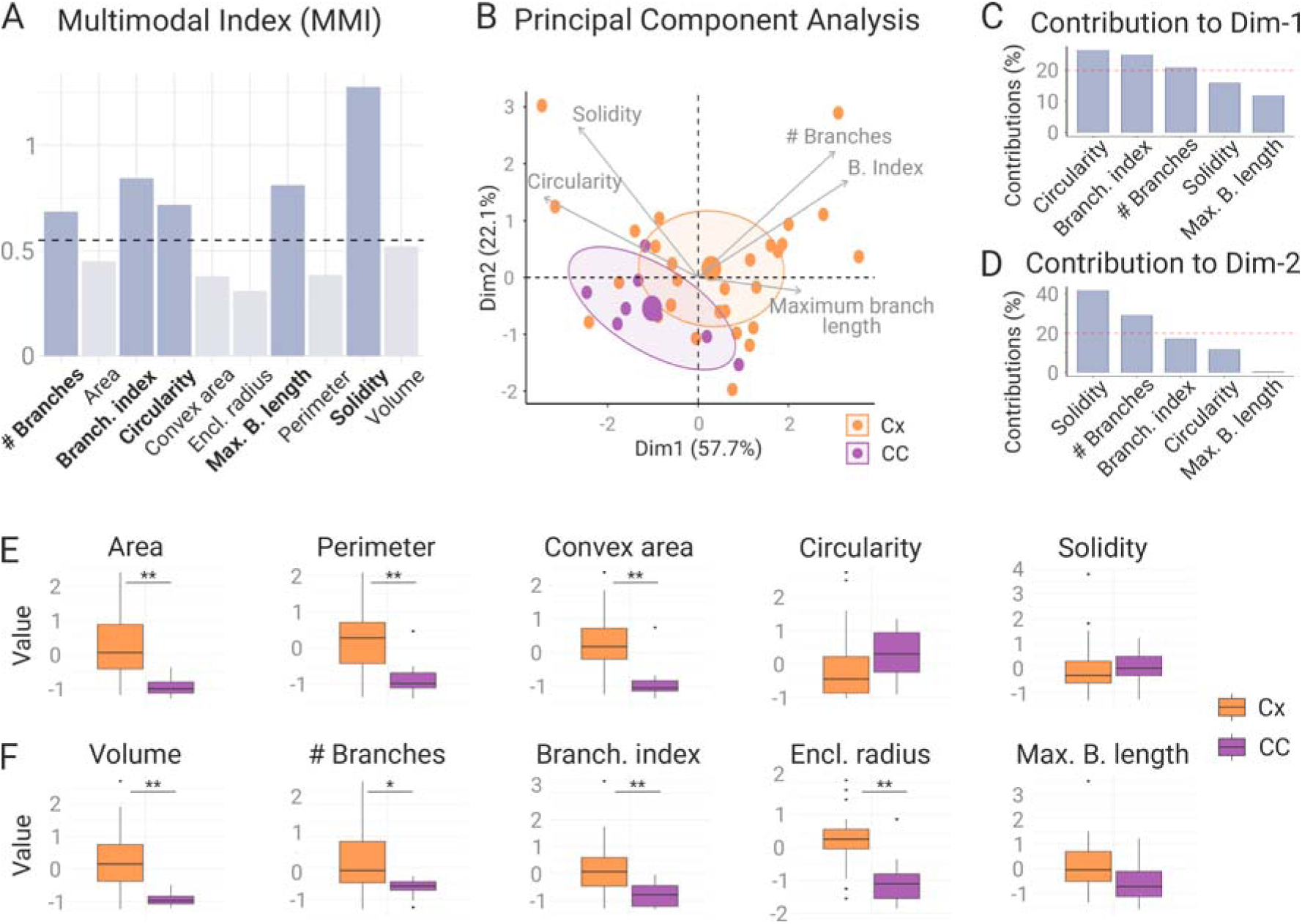
Regional Morphological Differences of NG2-Glia in the Cortex and Corpus Callosum. A) Multimodal Index (MMI) analysis identified the most informative parameters for clustering: circularity, solidity, branch number, branching index, and maximum branch length (MMI > 0.55). (B) Principal Component Analysis (PCA) of NG2-glia from cortex (Cx, orange) and corpus callosum (CC, purple) revealed distinct clusters, with PC1 (57.7% variance) driven by circularity, branching index, and branch number, and PC2 (22.1% variance) by solidity and branch number. (C, D) Bar plots showing the contribution of each parameter to PC1 (C) and PC2 (D). (E) Bar plots comparing size descriptors (area, perimeter, convex hull area, volume, enclosing radius) between NG2-glia from cortex and corpus callosum. (F) Bar plots comparing shape descriptors (branch number, branching index, circularity, solidity, maximum branch length). Asterisks indicate statistical significance: * p < 0.05, ** p < 0.01.

We concentrated on the first two principal components, which together explained 79.80% of the total variance (Figure 3B-D). The first component accounted for 57.7% of the variance and was mainly influenced by circularity, branching index, and the number of branches (Figure 3B, C). In comparison, the second component, explaining 22.1% of the variance, was primarily shaped by solidity and the number of branches (Figure 3B, D).

This analysis showed a degree of separation between cells from the two regions. NG2-glia from the corpus callosum formed a distinct cluster from those in the cortex, though some overlap was still observed (Figure 3B). Interestingly, among the MMI-selected parameters, only the number of branches and branching index accounted for the observed differences between NG2-glia in white matter and gray matter. No significant regional differences were found in circularity, solidity, or maximum branch length (Figure 3C, D).

Direct comparison of morphological features between cortical and corpus callosum NG2-glia revealed that circularity, solidity (Figure 3E), and maximum branch length (Figure 3F) were consistent across regions. Conversely, both the number of branches and the branching index were significantly higher in the cortex compared to the corpus callosum (Figure 3F).

In addition to regional variations in the two shape-related parameters—branch number and branching index—NG2-glial cells also showed differences in size-related features depending on their location. Notably, NG2-glia located in the cortex had significantly greater area, perimeter, convex hull area (Figure 3E), volume, and enclosing radius (Figure 3F) than those found in the corpus callosum.

These findings indicate that NG2-glia display region-specific morphological characteristics when comparing cells in the cortex versus those in the corpus callosum. Specifically, NG2-glial cells in the cortex possess a morphometric signature marked by increased size and more elaborate processes compared to their counterparts in the corpus callosum.

### 3.4 Morphological Layering of NG2-Glia Across the Cortical Layers

Due to the notable variability in cortical cell distribution highlighted by the PCA analysis (Figure 3B), we aimed to investigate whether NG2-glia display morphological distinctions across different cortical layers. To delineate these layers, we used Cux1 and CTIP2 as reference markers: layer 1 (L1), layers 2-4 (L2-4) for the upper cortex, and layers 5-6 (L5-6) for the lower cortex. Additionally, CTIP2 was utilized to identify the corpus callosum (Figure 4A). StarTrack-labeled NG2-glia, identified by cytoplasmic labeling, were detected across all cortical layers as well as in the corpus callosum (Figure 4B). This assessment included the same population of NG2-glial cells used in the broader white versus gray matter comparison. We detected NG2-glia in layer 1 (n = 8 cells), layers 2-4 (n = 8 cells), layers 5-6 (n = 12 cells), and in the corpus callosum (n = 8 cells), as identified by the appropriate neuronal markers for each cortical layer.

**Figure 4.**
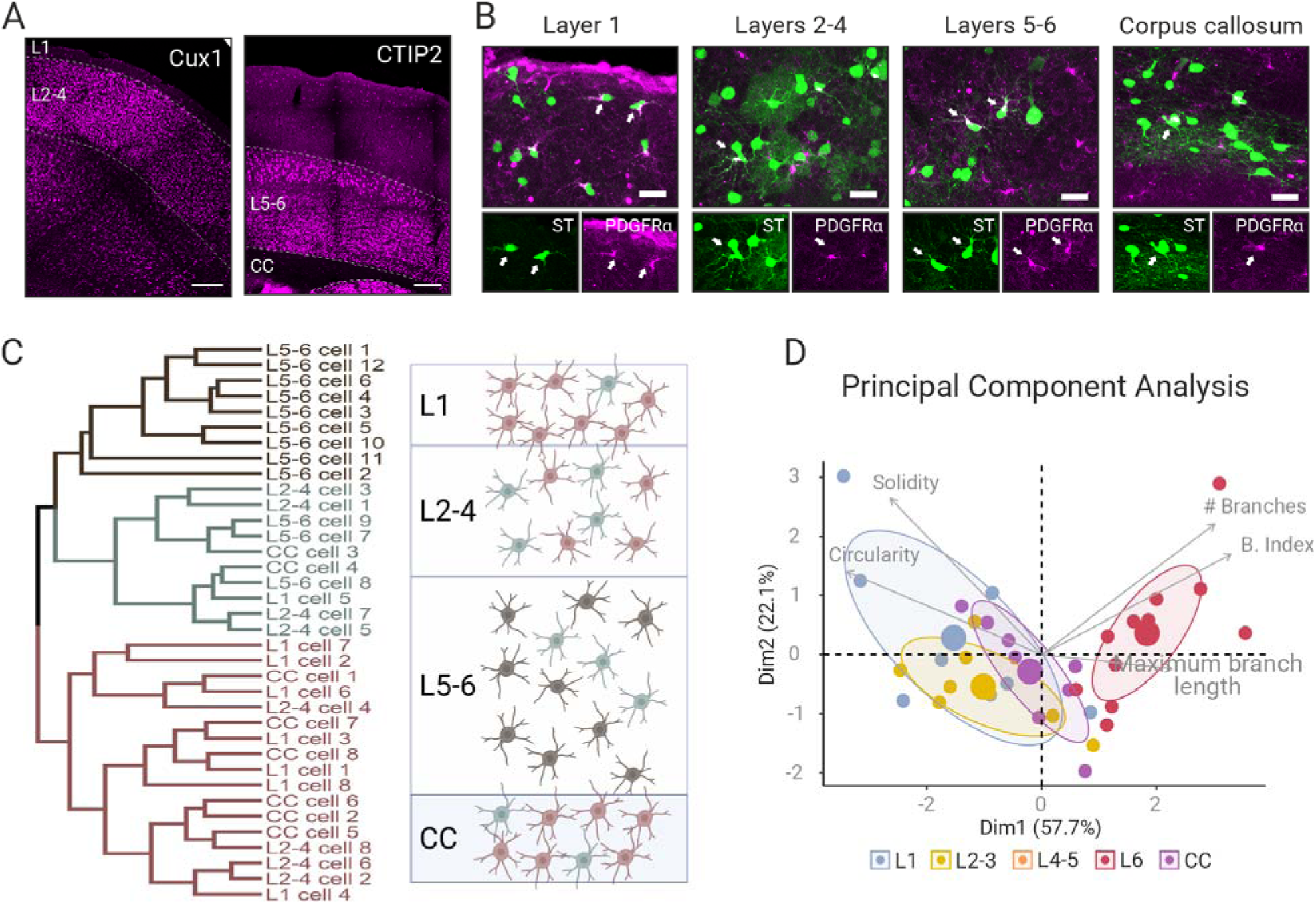
Morphological Variability of NG2-Glia Across Cortical Layers and the Corpus Callosum. (A) Cortical layers were delineated using Cux1 (L1, L2–4) and CTIP2 (L5–6, CC) immunolabeling. (B) Representative confocal images of StarTrack-labeled NG2-glia (green) co-labeled with PDGFRα (magenta) across layers. White arrows indicate double-labeled cells. Scale bars: 20 μm. (C) Hierarchical clustering of NG2-glia based on MMI-derived parameters revealed three main clusters: a compact cluster for L5–6 cells, a merged cluster for L1 and CC cells, and a dispersed cluster for L2–4 cells. Right: schematic of clustering patterns across regions. (D) PCA confirmed clustering patterns, showing that NG2-glia from L5–6 exhibited higher branch numbers and branching index, while those from L1 and CC displayed higher solidity and circularity.

To examine potential grouping patterns of NG2-glia based on their cortical layer location, we conducted hierarchical clustering of NG2-glial cells using the five morphometric parameters identified by MMI (circularity, solidity, branch number, branching index, and maximum branch length) (Figure 4C). Clustering was performed using complete linkage and Euclidean distance metrics. The resulting dendrogram revealed three main clusters, as depicted schematically (Figure 4C, right): 1) a compact cluster composed predominantly of NG2-glia from L5–6, suggesting strong morphological similarity within these deep cortical layers (Figure 4C, left, upper segment); 2) A merged cluster containing NG2-glia from L1 and the corpus callosum (Figure 4C, left, lower segment), indicating shared morphological features between these regions and 3) A more dispersed cluster formed by NG2-glia from L2–4, which were distributed across two separate branches of the dendrogram (Figure 4C, left, middle and lower segments), suggesting higher morphological heterogeneity within these mid-cortical layers.

These clustering patterns were further supported by PCA analysis (Figure 4D), aligning with the results obtained from hierarchical clustering (Figure 4C). PCA revealed that NG2-glia from L5–6 occupied a distinct morphological space, characterized by a greater number of branches and higher branching index. In contrast, NG2-glia from L1 and the corpus callosum clustered together, marked by higher solidity and circularity values /Fig. 4D) NG2-glia from L2–4 spanned an intermediate morphological profile, displaying greater heterogeneity and overlapping features with both deep and superficial cortical layers.

### 3.5 Regional Comparison of the Morphological Profiles of NG2-Glia

To comprehensively assess NG2-glial morphology across different regions, we performed a detailed analysis of size- and shape-related descriptors across cortical layers and the corpus callosum. These descriptors provided quantitative measures of NG2-glial cell structure, including area, perimeter, convex hull area, volume, enclosing radius, branch number, maximum branch length, branching index, solidity, and circularity. (Figure 5).

**Figure 5.**
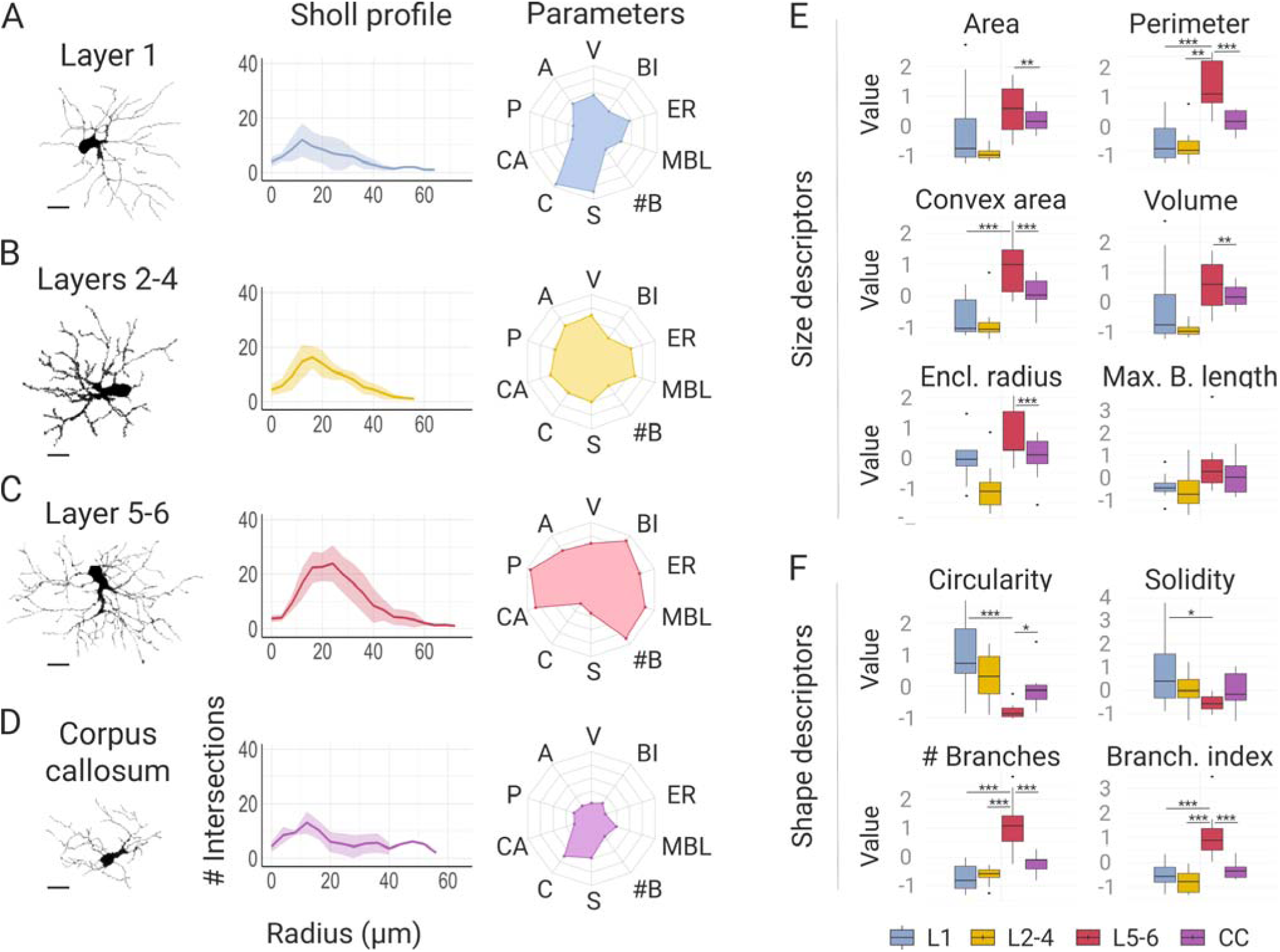
Morphological Profiles of NG2-Glia Across Cortical Layers and the Corpus Callosum. (A–D) Sholl analysis and radar plots of morphometric profiles for NG2-glia from (A) L1, (B) L2–4, (C) L5–6, and (D) CC. Radar plots include normalized metrics: volume (V), branching index (BI), enclosing radius (ER), maximum branch length (MBL), branch number (#B), solidity (S), circularity (C), convex hull area (CA), perimeter (P), and area (A). (E, F) Bar plots comparing size (E) and shape (F) parameters across regions. L5–6 cells displayed larger size and higher arbor complexity, while L1 and CC cells were more compact and spherical. Asterisks indicate significance: * p < 0.05, ** p < 0.01, *** p < 0.001.

Sholl analysis across layers revealed that NG2-glia located in layers 5–6 exhibited a greater number of intersections and extended to larger enclosing radii (Figure 5C) compared to those in layer 1, layers 2–4, or the corpus callosum (Figure 5A, B, D). This finding suggests that NG2-glia in the deeper cortical layers possess more elaborate and far-reaching arbors than in the upper layers or corpus callosum.

When comparing the normalized morphometric profiles of NG2-glia across layers (radar plots in Figure 5A–D, right panels), we observed that cells in L5–6 were consistently larger and exhibited more complex branching patterns, while those in L1 and the corpus callosum showed more compact and simpler morphologies. (Figure 5A, D). Cells in L2–4 exhibited intermediate traits (Figure 5B) bridging the morphological profiles of superficial and deep cortical layers (Figure 5C).

Quantitative comparisons in size (Figure 5E) and shape (Figure 5F) parameters revealed that NG2-glia in L5–6 had significantly greater area, perimeter, convex hull area, volume, and enclosing radius compared to cells in L1 and the corpus callosum. Although NG2-glia in L5–6 also showed higher maximum branch length values, these differences did not reach statistical significance (Figure 5E).

Regarding shape descriptors, NG2-glia in layers 5–6 exhibited higher branch numbers and branching indices, indicating increased arbor complexity. In contrast, NG2-glia in L1 and the corpus callosum displayed higher solidity and circularity values, reflecting a more compact, spherical morphology. NG2-glia in L2–4 showed intermediate values, consistent with their mixed morphological profile. (Figure 5F).

Together, these findings indicate that NG2-glia exhibit **region-specific morphologies** across the cortex and corpus callosum, with cells in deep cortical layers (L5–6) forming a morphometrically distinct population characterized by larger size and greater process complexity, while cells in superficial layers and white matter display more compact and less branched profiles.

### 3.6 Morphological Comparison between Astrocytes and NG2-Glia

To further contextualize the morphological features of NG2-glia, we compared them with astrocytes from the same cortical layers and the corpus callosum, using a dataset (n=45 cells) obtained from a previous study that employed the same StarTrack-based reconstruction methodology (Delgado-García et al., 2024). PCA analysis revealed clear morphological separation between astrocytes and NG2-glia across all examined regions: upper cortical layers (Figure 6A), lower cortical layers (Figure 6B), and the corpus callosum (Figure 6C).).

**Figure 6.**
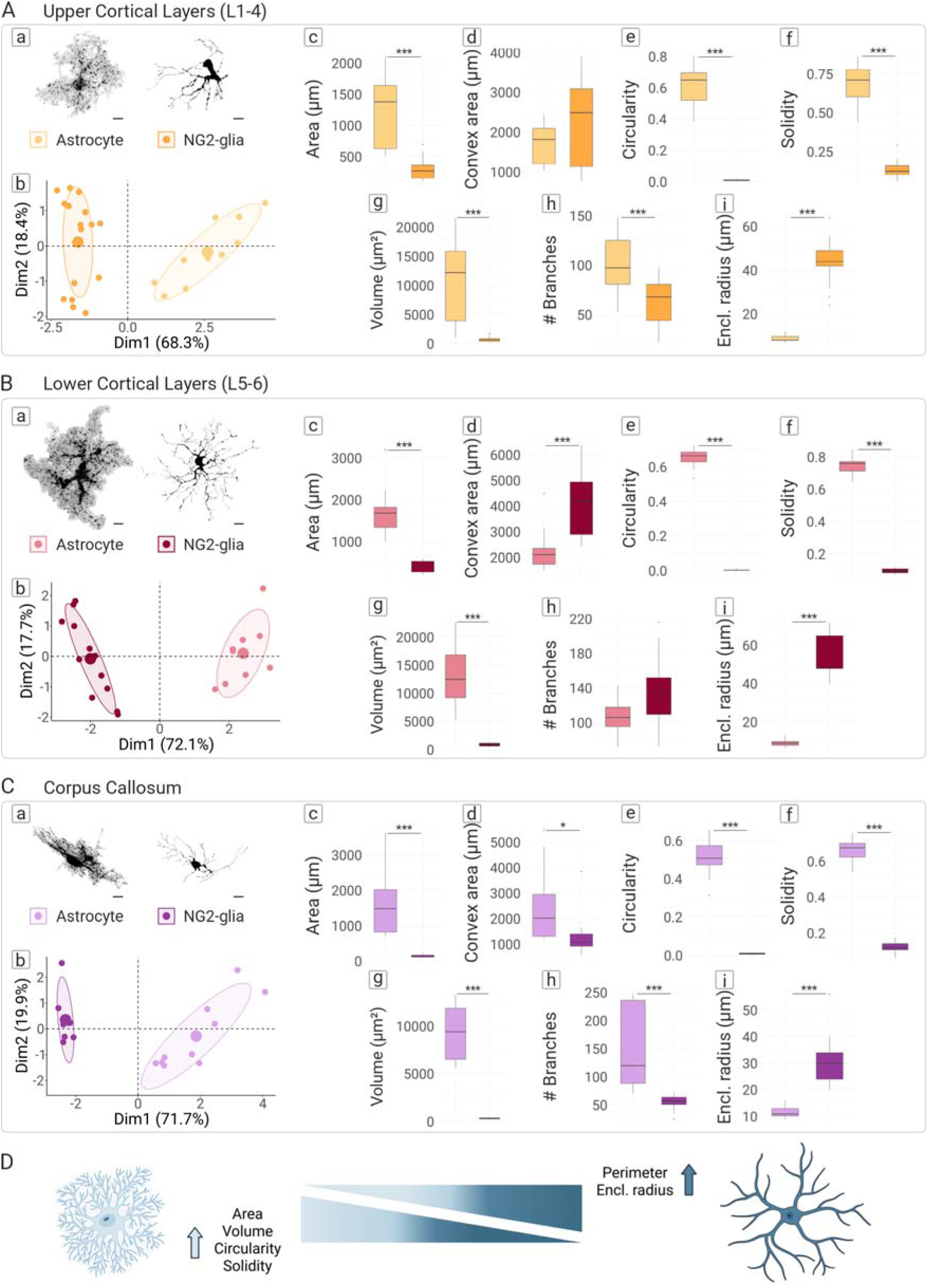
Morphological differences between astrocytes and NG2-glial cells in different cortical layers. (A–C) Comparisons of astrocytes and NG2-glia in (A) upper cortical layers (L1–4), (B) lower cortical layers (L5–6), and (C) corpus callosum (CC). Each panel includes: (a) representative reconstructions, (b) PCA plots showing clear separation of astrocytes (blue) and NG2-glia (green), and (c–j) box plots comparing area, perimeter, convex hull area, volume, maximum branch length, circularity, solidity, branch number, and enclosing radius. (D) Summary schematic: astrocytes (left) exhibit higher area, volume, circularity, and solidity; NG2-glia (right) display greater perimeter and enclosing radius. Asterisks indicate significance: * p < 0.05, ** p < 0.01.

Comparative analyses demonstrated that astrocytes exhibited significantly greater area and volume than NG2-glia in all regions examined (Figures 6A.c, B.c, C.c, and 6A.h, B.h, C.h, respectively). Conversely, NG2-glia showed higher perimeter and enclosing radius values (Figures 6A.d, B.d, C.d, and 6A.j, B.j, C.j), suggesting that although astrocytes occupy more volume, NG2-glia extend their processes further. NG2-glia exhibited lower circularity and solidity across all regions (Figures 6A.f, B.f, C.f, and 6A.g, B.g, C.g), indicating a less compact morphology compared to astrocytes. Moreover, astrocytes displayed higher branch numbers in both upper cortical layers and the corpus callosum (Figures 6A.i, C.i), reflecting greater process complexity Interestingly, convex hull area comparisons revealed that NG2-glia had higher convex hull areas than astrocytes in the lower cortical layers (Figure 6B.e), but lower values in the corpus callosum (Figure 6C.e).

These findings suggest that while astrocytes occupy a more compact, volumetrically dense domain (Figure 6D, left), NG2-glia extend longer processes that cover broader territories, albeit with lower process density (Figure 6D, right).

Collectively, our results provide a comprehensive, quantitative characterization of NG2-glia morphology across cortical layers and the corpus callosum. NG2-glia in deep cortical layers (L5–6) exhibit a distinct morphological signature characterized by larger size and greater process complexity, while those in superficial layers (L1) and white matter (corpus callosum) display a more compact, spherical morphology. NG2-glia in mid-cortical layers (L2–4) demonstrate intermediate features, bridging the profiles of deep and superficial populations. Moreover, comparative analyses reveal that NG2-glia and astrocytes exhibit fundamentally distinct morphologies: astrocytes form denser, more compact arbors, while NG2-glia extend thinner, longer processes across broader spatial domains. These structural differences likely reflect their distinct functional roles within neural circuits.

## 4 Discussion

NG2-glia constituted a specialized glial population distinct from astrocytes, oligodendrocytes, and microglia, demonstrating progenitor potential, synaptic connectivity, and environmental responsiveness (Richardson et al., 2011; Dimou and Götz, 2014; Dimou and Simons, 2017; Jäkel and Dimou, 2017).

Our findings highlight the morphological diversity of NG2-glia and support the growing understanding of their functional roles (Auguste et al., 2022; Bergles et al., 2000; Buchanan et al., 2022; Dimou et al., 2008; Mount et al., 2019; Seo et al., 2013; Serwanski et al., 2017; Wigley & Butt, 2009; Xiao et al., 2022; Young et al., 2013; Zhang et al., 2021a). Our data reveal rich morphological profiles as described in other brain regions (Hill et al., 2013; Viganò et al., 2013; Young et al., 2013).

We reveal clear morphological differences between NG2-glia in gray matter (cortex) and white matter (corpus callosum). This is consistent with the known functional heterogeneity of NG2-glia across these regions, both in terms of proliferation rates and their potential to differentiate into other cell types (Hill et al., 2013; Dimou and Gallo, 2015). The smaller size and reduced branching complexity of NG2-glia in the corpus callosum may be linked to a lower basal proliferation rate and a higher tendency to differentiate into oligodendrocytes (Psachoulia et al., 2009; Simon et al., 2011; Viganò et al., 2013; Young et al., 2013; Lodato and Arlotta, 2015). Additionally, this morphological phenotype may reflect their greater contact with nodes of Ranvier in white matter compared to their counterparts in the cortex (Serwanski et al., 2017).

Within the cortex, deep-layer NG2-glia (L5–6) displayed enlarged somata, extensive arborizations, and higher branching indices compared to superficial layers (L1–4) and the corpus callosum. Interestingly, this laminar pattern mirrors neuronal organization: deep layers contain projection neurons targeting subcortical structures and the thalamus, while superficial layers are home to callosal projection neurons, intracortical processing neurons, and interneurons (Lodato and Arlotta, 2015). The alignment between NG2-glial layering and neuronal subtypes organization suggests that synaptic interactions between NG2-glia and neurons may vary depending on the type of neuron involved. This supports the idea that NG2-glia perform diverse, context-dependent functions shaped by the specific neuronal environment in which they reside. In fact, these synaptic interactions with neurons are a unique feature of NG2-glia (Bergles et al., 2000), a property that sets them apart from other glial cell types. Although this phenomenon is not yet fully understood, NG2-glia are known to receive both glutamatergic and GABAergic inputs from neurons (Ziskin et al., 2007; Vélez-Fort et al., 2010; Orduz et al., 2015). These cells express a variety of neurotransmitter receptors (Larson et al., 2016), along with ion channels that allow them to respond to synaptic signals by altering their membrane potential during transmission (Wigley and Butt, 2009).

Remarkably, NG2-glia in layer 1 and the corpus callosum share similar morphological features despite being located in separated anatomical regions. This suggests that local microenvironmental cues, rather than mere anatomical proximity, play a key role in shaping their structure. In addition, we observed considerable morphological variability within the upper cortical layers, particularly layers 2-4, pointing to the presence of distinct NG2-glial subpopulations within the same region. This highlights the complexity of NG2-glial identity and the importance of studying these cells in finer spatial resolution.

Thus, the ability of NG2-glia to integrate synaptic and electrophysiological inputs, combined with their intrinsic molecular plasticity, underscores their essential role in both the development of neural circuits and the maintenance of brain homeostasis.

The layering of NG2-glia throughout the cortex, mirroring the laminar structure of neurons, further supports the idea that NG2-glia, like astrocytes (Lanjakornsiripan et al., 2018; Xin Tan et al., 2023; Baldwin et al., 2024) adapt their shape, and likely their function, in response to the demands of their local neuronal environment. Our findings reinforce the strong link between structure and function in NG2-glia, suggesting that their morphology is not just a passive reflection of their location, but an active adaptation for specialized roles within neural local circuits. In line with this idea, recent work has shown that locomotion-responsive NG2-glia of the somatosensory cortex simplify their dendritic arbor while increasing soma size, whereas non-responsive cells retain more elaborate and stable morphologies (Fiore et al., 2023). This functional dichotomy further supports the notion that NG2-glia exhibit structural plasticity that may reflect underlying differences in functional states, a concept that is consistent with the regional and laminar morphological specializations we observed in our study.

Beyond regional differences, our study also highlights key structural differences between NG2-glial cells and astrocytes. Astrocytes, known for regulating synaptic activity through tripartite synapses (Araque et al., 1999), have denser and more compact arbors, occupying larger volumes with higher circularity and solidity. NG2-glia, in contrast, have fewer branches and smaller somas, but extend longer processes and occupy larger territories than astrocytes, despite having a smaller volume. This may allow NG2-glia to influence broader areas and interact with more neurons.

These morphological differences, together with the ability of NG2-glia to receive direct synaptic input from neurons (Ge et al., 2006; Ziskin et al., 2007; Vélez-Fort et al., 2010; Orduz et al., 2015; Mount et al., 2019), supports the idea that their morphology is specifically adapted for a unique, integrative role within neural circuits.

Our findings suggest that astrocytes and NG2-glia use different but complementary structural strategies: astrocytes form dense, highly branched networks optimized for local synaptic modulation, while NG2-glia extend longer, sparser processes that may help them interact across wider areas and monitor neuronal activity more broadly.

This study also highlights the strength of the StarTrack system for studying glial cell morphology. By labeling the cytoplasm of individual cells, StarTrack provides high-resolution detail that makes it possible to reconstruct fine cellular processes and identify distinct morphological subtypes within diverse glial populations. Previous studies have demonstrated the versatility of StarTrack in lineage tracing, developmental studies, and injury paradigms (García-Marqués and López-Mascaraque, 2013; Martín-López et al., 2013; Bribian et al., 2018; Barriola et al., 2020; Delgado-García et al., 2024; Ojalvo-Sanz et al., 2025).

In conclusion, our work shows that NG2-glial morphology is not uniform, but rather exhibits region- and layer-specific signatures that likely reflect functional specialization. The distinct morphologies of NG2-glia across cortical layers and the corpus callosum, along with their clear divergence from astrocytes, highlight the importance of studying glial cells in their exact anatomical and functional settings. Future studies explore how developmental origins, regional signals, and functional demands interact to shape the diverse morphologies and roles of NG2-glia in the adult CNS. The combination of lineage tracing, functional imaging, and transcriptomics will help us better understand how NG2-glia adapt to the complex needs of neural circuits.

## 6 Author Contributions

LLM and SB contributed to the conception and design of the study. SB, CSP, ACOS, MFO and LMDG performed the methodological strategies. SB organized the database and performed the statistical analysis. SB and LLM wrote the first draft of the manuscript. All authors contributed to the manuscript revision, read, and approved the submitted version.

## Acknowledgments

We are very pleased to the Animal Facilities of the Instituto Cajal for the assistance. In addition, we are thankful to our lab.

## Ethics approval statement

Animal procedures were conducted in accordance with the European Directive 2010/63/EU and the regulations established by the Spanish Animal Care and Committee. Experimental protocols received approval from the Bioethics Committee of Community of Madrid (Reference: PROEX 314/19)

